# Quantifying the polygenic architecture of the human cerebral cortex: Extensive genetic overlap between cortical thickness and surface area

**DOI:** 10.1101/868307

**Authors:** Dennis van der Meer, Oleksandr Frei, Tobias Kaufmann, Chi-Hua Chen, Wesley K. Thompson, Kevin S. O’Connell, Jennifer Monereo Sánchez, David E.J. Linden, Lars T. Westlye, Anders M. Dale, Ole A. Andreassen

**Author notes:** Corresponding authors &. Address: Kirkeveien 166, 0450 Oslo, Norway.

## Abstract

**Introduction:** The thickness of the cerebral cortical sheet and its surface area are highly heritable traits thought to have largely distinct polygenic architectures. Despite large-scale efforts, the majority of their genetic determinants remains unknown. Our ability to identify causal genetic variants can be improved by employing better delineated, less noisy brain measures that better map onto the biology we seek to understand. Such measures may have fewer variants but with larger effects, i.e. lower polygenicity and higher discoverability.

**Methods:** Using Gaussian mixture modeling, we estimated the number of causal variants shared between mean cortical thickness and total surface area. We further determined the polygenicity and discoverability of regional cortical measures from five often-employed parcellation schemes. We made use of UK Biobank data from 31,312 healthy White European individuals (mean age 55.5, standard deviation (SD) 7.4, 52.1% female).

**Results:** Contrary to previous reports, we found large genetic overlap between total surface area and mean thickness, sharing 4427 out of 7150 causal variants. Regional surface area was more discoverable (p=4.1×10^−6^) and less polygenic (p=.007) than regional thickness measures. We further found that genetically-informed and less granular parcellation schemes had highest discoverability, with no differences in polygenicity.

**Conclusions:** These findings may serve as a roadmap for improved future GWAS studies; Knowledge of which measures or parcellations are most discoverable, as well as the genetic overlap between these measures, may be used to boost identification of genetic predictors and thereby gain a better understanding of brain morphology.

---

The morphology of the human cerebral cortex is highly heritable, and identifying the genetic variants involved will have fundamental impact on our understanding of brain development. Despite large-scale efforts, the majority of these genetic variants remains unknown^1^. This is in part due to the genetic signal of cortical morphology being distributed across many causal variants, each having a small effect^2,3^. Our ability to identify causal SNPs can be improved not only by increasing sample sizes to boost statistical power, but also by employing better delineated, less noisy brain measures that better map onto the biology we seek to understand. Such measures might have fewer causal variants but with larger effects, i.e. lower polygenicity and higher discoverability. Quantifying these characteristics of the polygenic architecture across often-used cortical measures may therefore optimize the selection of the most informative measures, enhancing discovery.

The thickness of the cerebral cortical sheet and its surface area are two separable morphological measures which have been reported to follow differing trajectories over the lifespan^4^ and to be differentially associated with cognitive ability^5^.

They have further been reported to have largely distinct genetic determinants, suggesting the genetic underpinnings of these two measures should be assessed separately^6,7^. This claim is, however, based on estimates of genetic correlation, a genome-wide measure of the correlation of genetic effects on the two traits. Complex traits such as brain measures may have a substantial number of shared genetic loci^8^, even in the absence of genetic correlation, due to mixed directions of effects^9^. Identification of the fraction of shared causal variants between two brain phenotypes^9^, beyond genetic correlation, is valuable for an understanding of their biological relation. This overlap may further be exploited to boost identification of genetic factors for these traits^10^.

Here, we quantified important characteristics of the polygenic architecture of the cerebral cortex, estimating the genetic overlap between mean cortical thickness and total surface area, as well as the polygenicity, discoverability, and heritability of regional measures. This was achieved through Gaussian mixture modeling, using the MiXer tool^3,9^. We processed T1-weighted magnetic resonance image (MRI) scans from 31,312 healthy individuals with White European ancestry (mean age 55.5, standard deviation (SD) 7.4, 52.1% female), as part of the UK Biobank^11,12^, using the standardized pipeline of FreeSurfer. In addition to estimating the total surface area and mean cortical thickness, we divided each hemisphere into 34 regions using the Desikan-Killiany atlas with boundaries based on gyral and sulcal patterns^13^, extracting area and thickness estimates for each. For all measures, we regressed out age, sex, scanning site, a proxy of scan quality (FreeSurfer’s Euler number)^14^, and the first twenty genetic principal components. For the regional measures of thickness and area, we also regressed out the corresponding hemisphere-specific global measure, in accordance with previous work^1^. Making use of the UKB v3 imputed data, we performed genome-wide association studies (GWAS) on the pre-residualized measures through PLINK 2.0^15^. We then applied MiXeR to the resulting summary statistics, calculating polygenicity (estimated number of causal variants, NC), discoverability (proportion of phenotypic variance explained on average by a causal variant, 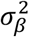), and narrow-sense heritability (the product of polygenicity and discoverability, i.e. proportion of phenotypic variance explained, h^2^). We excluded from our analyses regions where the ratio between the estimated heritability and its standard error (SE) was less than 3, as this suggests there may be insufficient signal in the data to reliably estimate MiXer parameters^9^. For more details on data processing, exclusion, and analysis, please see the Online Methods.

We identified a large degree of genetic overlap between total surface area and mean thickness, with a Dice coefficient of 0.76, see Figure 1A. This is in contrast to the negative genetic correlation as estimated through linkage-disequilibrium score regression (LDSC)^16^, displayed below the Venn diagram, which suggests a smaller degree of genetic overlap. The bivariate density plot, Figure 1B, illustrates mixed directions of effects for many SNPs, which explains these apparent conflicting findings; some SNPs have the same direction of effect on both traits, while others have a positive effect on area and a negative effect on thickness or vice versa, with the net result being a smaller negative correlation.

**Figure 1.**
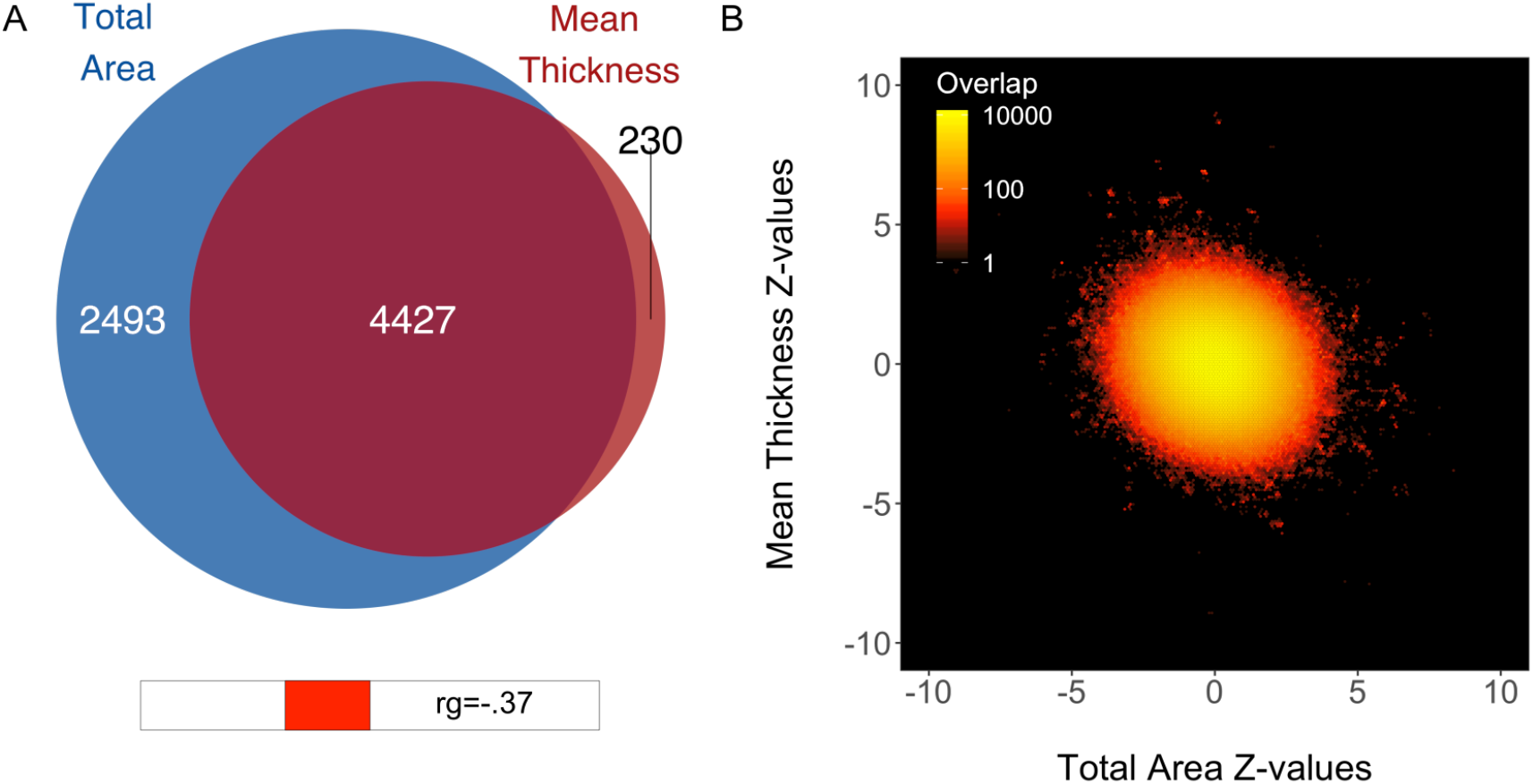
Genetic overlap of total surface area and mean thickness. **A)** Venn diagram depicting the estimated number of causal variants shared between total surface area and mean thickness and unique to either of them. Below the diagram, we show the estimated genetic correlation. B) Bivariate density plot, illustrating the relationship between the observed GWAS Z-values for total area (on the x-axis) and mean thickness (on the y-axis).

Our analysis revealed that total surface area is more heritable (h^2^=.31, SE=.02) than mean thickness (h^2^=.23, SE=.02). in accordance with previous findings^1^. We show that surface area has a marginally higher polygenicity (NC=6920, SE=1278) than mean thickness (NC=4657, SE=1114), at similar levels of discoverability (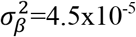, SE=6.5×10^−6^ vs 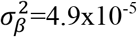, SE=8.9×10^−6^).

Using the Desikan-Killiany regional estimates, we observed a strong trade-off between the polygenicity and discoverability (Spearman’s r_s_=−.88, p<1×10^−16^), as shown in Figure 2A. Regional heritability was positively correlated with discoverability (r_s_=.37, p=4.4×10^−5^), but not with polygenicity (r_s_=−.08, p=.37). Regional area was significantly less polygenic, and more discoverable and heritable than regional thickness, as shown in Figure 2B-D. Further, we found that the size of the brain regions was positively correlated with their discoverability (r_s_=.41, p=.001) and heritability (r_s_=.38, p=.002), and negatively related to their polygenicity (r_s_=−.27, p=.047), for the area-specific estimates only. For the full results, and an overview of parameter estimates per region, see the Extended Data.

**Figure 2.**
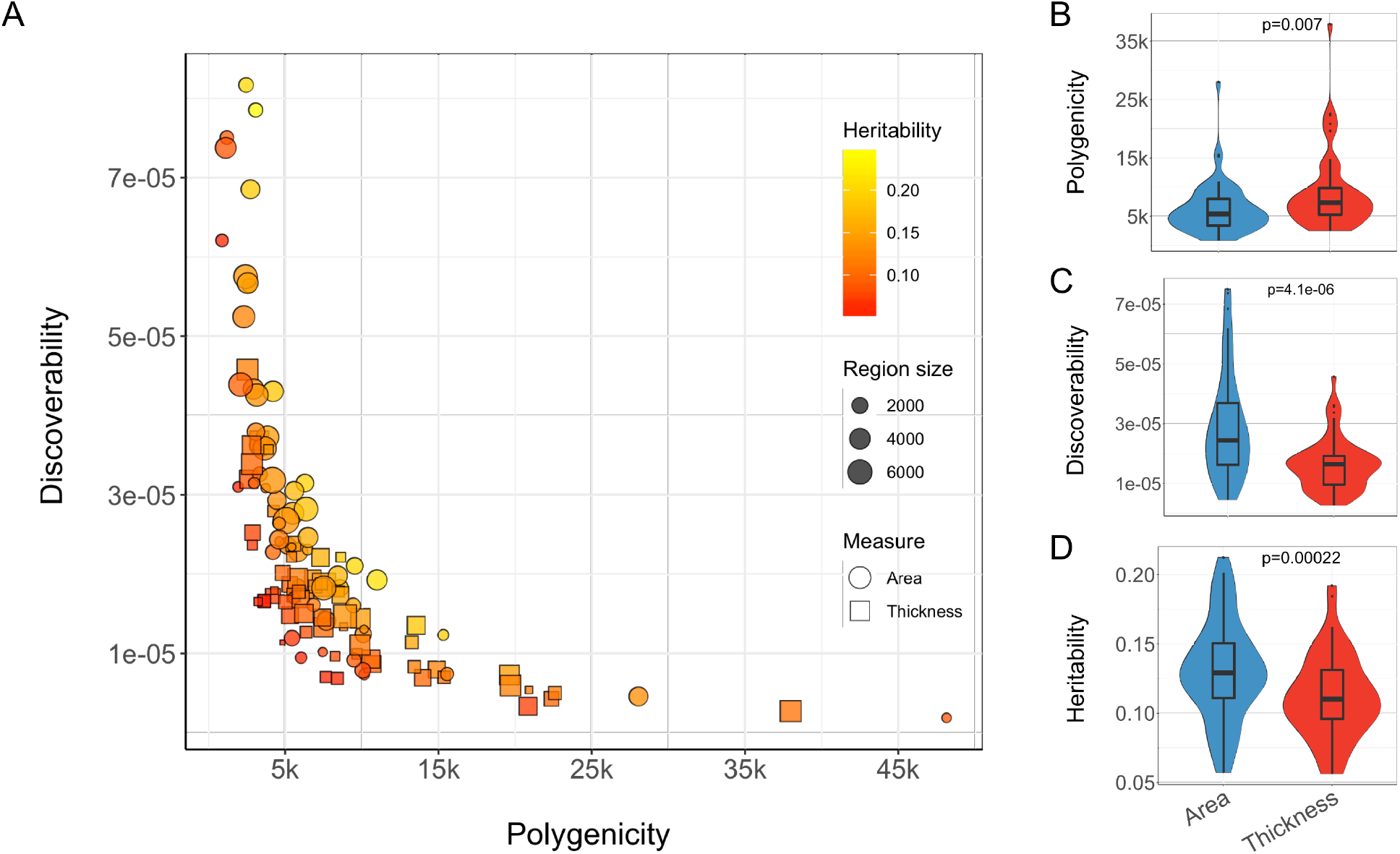
Polygenicity, discoverability and heritability of regional cortical surface area and thickness. A) Scatter plot depicting the relationship between estimates of polygenicity (x-axis) and discoverability (y-axis) for regional cortical measures. The shape of the data points distinguishes between surface area and thickness estimates, the point size reflects the size of the region, and the color relates to the estimated heritability. B-D) Violin plots comparing the polygenicity, discoverability and heritability (on the y-axis) of regional area and thickness estimates (x-axis). Significance, indicated at the top of each graph, is calculated through the Wilcoxon signed-rank test.

Given that the definition of regions likely plays an important role in the results of imaging genetics studies, we additionally compared the Desikan-Killiany atlas with three alternative popular parcellation schemes: 1) the Chen *et al*. atlas, dividing each hemisphere into 12 clusters based on genetic correlations of surface area from a large twin study^17^, 2) the Yeo *et al*. atlas, which comes in either a 7 or 17-cluster solution based on resting-state functional connectivity^18^, and 3) the Glasser *et al*. atlas, with the boundaries of a 180 regions per hemisphere formed based on multimodal imaging data^19^. Thus these schemes differ both in the biological basis of their parcellation, and their granularity. Of the five schemes, for the area measures, the Chen *et al*. parcellation was both the most discoverable and the most heritable. There were no significant differences in polygenicity between the parcellation schemes. These results are summarized in Figure 3. There were no differences in discoverability of regional thickness between the atlases (see Extended Data).

**Figure 3.**
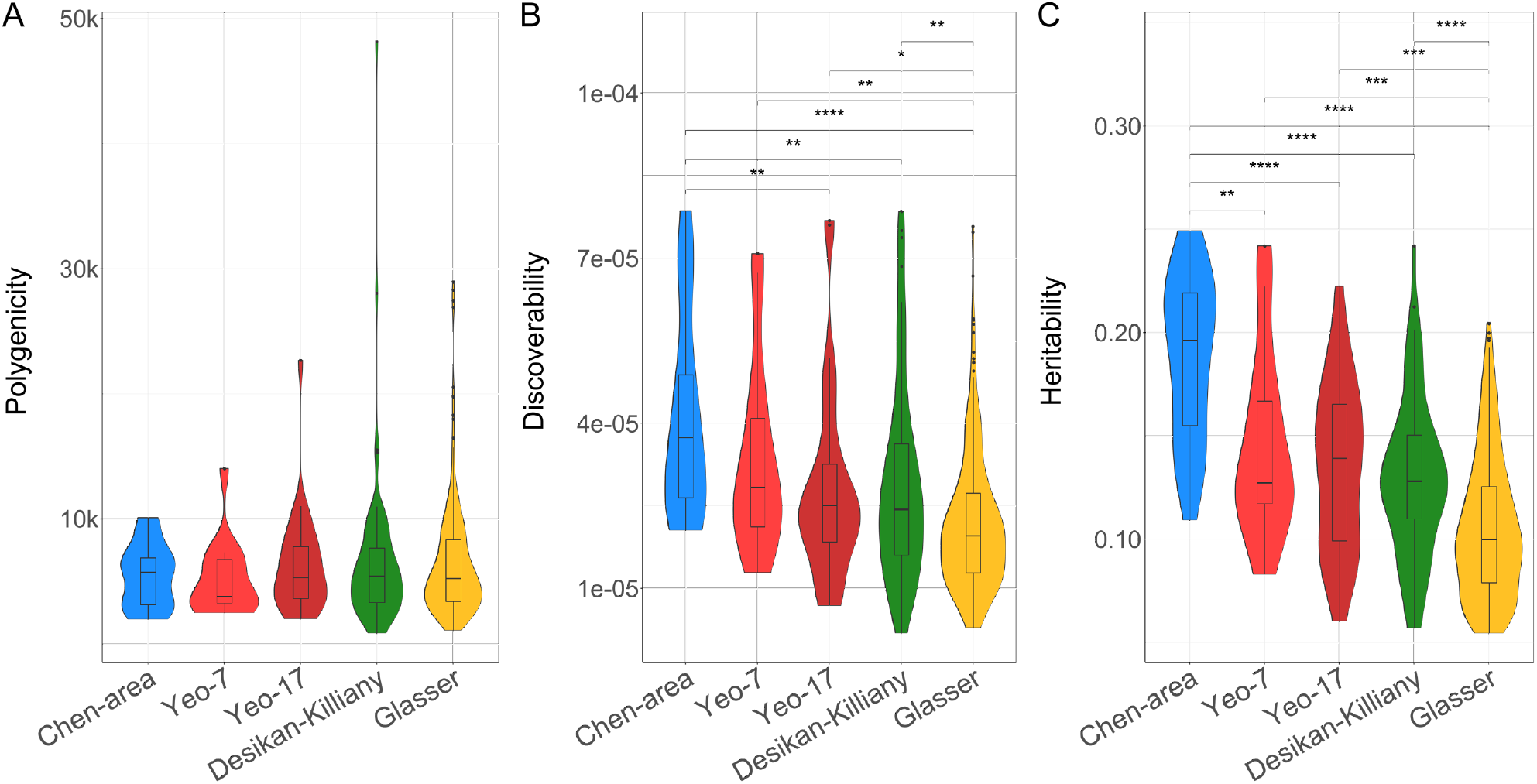
Comparisons of discoverability and polygenicity of regional surface area across parcellation schemes. Violin plots comparing the polygenicity (A), discoverability (B) and heritability (C) (on the y-axis) of the five different parcellation schemes (x-axis). Significance of each paired comparison, indicated at the top of each graph, is calculated through the Wilcoxon signed-rank test with *:p<.05, **:p<.01, ***:p<.001, and ****:p<.00001.

These findings provide new insights into the polygenic architecture of the cerebral cortex, with three main take-home messages. First, there is large genetic overlap between total surface area and mean thickness. Second, regional surface area measures are more discoverable and less polygenic than regional thickness measures. And third, the genetically-informed and less granular parcellation schemes had highest discoverability.

We provide evidence of strong genetic overlap between total surface area and mean thickness, contrary to previous reports based on genetic correlations^6,7^. The known, phenotypic and genetic, weak negative correlation therefore appears not to be the result of a moderate amount of shared genetic variants with opposing direction of effects, but rather a large amount of shared genetic variants with a mixture of opposite and same direction of effects. This adds to recent literature revealing high levels of pleiotropy between brain-related behavioral traits and disorders^20^. For instance, nearly all causal variants for schizophrenia are shared with educational attainment despite a negligible genetic correlation^9^, in accordance with research identifying many shared genetic loci between the two traits without a clear pattern of sign concordance^21,22^. This is critical for our understanding of the biological relationships between brain-related traits, and how to study them.

We further found that regional area is more heritable than regional thickness, in line with previous reports^23^, and we show that this is driven by higher discoverability. This matches findings from a recent GWAS study, with 187 genome-wide significant hits for regional area versus only 12 for regional thickness^1^. There appears to be more variation in the distribution of the estimates for regional surface area compared to regional cortical thickness, suggesting the latter has a more homogeneous polygenic architecture across regions. Larger differences between the polygenic architectures of regional surface area measures may also underlie the somewhat higher polygenicity for total surface area compared to mean thickness. It may further explain why most genome-wide significant variants are found for surface area, as only those regions with high discoverability have effects that may be identified with current sample sizes.

Our comparison of different parcellation schemes indicates that the choice of scheme makes a significant difference on the outcome of imaging genetics studies. While all schemes had similar levels of polygenicity, the Chen *et al*. parcellation performed best in terms of discoverability. Besides the fact that this parcellation was based on twin data, i.e. genetically informed, granularity may play an important role in the performance of the schemes we compared; measurement noise due to inaccuracy of boundary placement will disproportionally affect smaller regions, thereby lowering heritability^23,24^ and discoverability, compounding the multiple comparisons problem faced by studying more granular schemes. Boundary placement accuracy at the individual level is of less importance for thickness estimates^23^, likely contributing to why we only saw differences between the schemes for the area estimates.

To conclude, we revealed that surface area and thickness share a considerable number of genetic variants and provide the first estimates of discoverability and polygenicity of regional cortical measures across parcellation schemes. These findings may serve as a roadmap for improving future studies. Knowledge of which measures or parcellations are most discoverable, and why, as well as the genetic overlap between these measures, can be exploited to boost identification of genetic predictors and thereby gain a better understanding of brain morphology.

## Author contributions

D.v.d.M., O.F., A.M.D. and O.A.A. conceived the study; D.v.d.M. and T.K. pre-processed the data; D.v.d.M. and O.F. performed all analyses, with conceptual input from T.K., A.M.D. and O.A.A.; All authors contributed to interpretation of results; D.v.d.M. drafted the manuscript and all authors contributed to and approved the final manuscript.

## Materials & Correspondence

The data incorporated in this work were gathered from public resources. The code is available via https://github.com/precimed/mixer (GPLv3 license) upon publication of this study. Correspondence and requests for materials should be addressed to

## Acknowledgements

The authors were funded by the Research Council of Norway (276082, 213837, 223273, 204966/F20, 229129, 249795/F20, 225989, 248778, 249795), the South-Eastern Norway Regional Health Authority (2013-123, 2014-097, 2015-073, 2016-064, 2017-004), Stiftelsen Kristian Gerhard Jebsen (SKGJ-Med-008), The European Research Council (ERC) under the European Union’s Horizon 2020 research and innovation programme (ERC Starting Grant, Grant agreement No. 802998) and National Institutes of Health (R01MH118281, R01GM104400). This work was partly performed on the TSD (Tjeneste for Sensitive Data) facilities, owned by the University of Oslo, operated and developed by the TSD service group at the University of Oslo, IT-Department (USIT). Computations were also performed on resources provided by UNINETT Sigma2 - the National Infrastructure for High Performance Computing and Data Storage in Norway.

## Online Methods

### Sample

We made use of data from participants of the UKB population cohort, obtained from the data repository under accession number 27412. The composition, set-up, and data gathering protocols of the UKB have been extensively described elsewhere^11^. For this study, we selected White Europeans that had undergone the neuroimaging protocol. We excluded 1286 individuals with a primary or secondary ICD10 diagnosis of a neurological or mental disorder, as well as 691 individuals with bad structural scan quality as indicated by an age and sex-adjusted Euler number^14^ more than three standard deviations lower than the scanner site mean. Our final sample size was n=31,312, with a mean age of 55.5 years (SD=7.4). 52.1% of the sample was female.

### Data preprocessing

T1 scans were collected from three scanning sites throughout the United Kingdom, all on identical Siemens Skyra 3T scanners with a 32-channel receive head coil. The UKB core neuroimaging team has published extensive information on the applied scanning protocols and procedures, which we refer to for more details^12^. The T1 scans were obtained from the UKB data repositories and stored locally at the secure computing cluster of the University of Oslo. We applied the standard “recon-all” processing pipeline of Freesurfer v5.3, performing automated surface-based morphometry segmentation^13^. From the output, we extracted global and regional estimates of cortical thickness and surface area. For the primary analyses, we made use of the Desikan-Killiany atlas, dividing each hemisphere into 34 regions, based on gyral and sulcal patterns^13^. We additionally extracted regional estimates of cortical thickness and surface area using three other parcellation approaches: 1) the Chen *et al*. surface area atlas, which divides each hemisphere into 12 clusters, based on a data-driven fuzzy clustering technique applied to estimates of genetic correlations derived from cortical surface area data from 406 monozygotic and dizygotic twins^17^. For this, we made use of the GCLUST phenotype extraction protocol (https://github.com/ENIGMA-git/GCLUST), 2) The Yeo *et al*. atlas, which provides a 7 and a 17 cluster solution of dividing each hemisphere, based on functional connectivity patterns in resting-state fMRI data of a 1000 subjects^18^ (https://surfer.nmr.mgh.harvard.edu/fswiki/CorticalParcellation_Yeo2011), and 3) the Glasser *et al*. atlas, dividing the hemispheres into 180 regions, based on multimodal MRI data from the Human Connectome Project (HCP) and an objective semi-automated neuroanatomical approach^19^ (https://figshare.com/articles/HCP-MMP1_0_projected_on_fsaverage/3498446).

We subsequently regressed out age, sex, scanner site, Euler number, and the first twenty genetic principal components from each measure. We further regressed out a hemisphere-specific global measure for each of the regional measures: mean thickness for the regional thickness measures, and total surface area for the regional surface area measures. Subsequently, we applied a rank-based inverse normal transformation to the residuals of each measure, ensuring normally distributed input into the GWAS.

### Univariate GWAS procedure

We made use of the UKB v3 imputed data, which has undergone extensive quality control procedures as described by the UKB genetics team^25^. After converting the BGEN format to PLINK binary format, we additionally carried out standard quality check procedures, including removal of SNPs with low imputation quality scores (INFO <.5), filtering out individuals with more than 10% missingness, SNPs with more than 5% missingness, and SNPs failing the Hardy-Weinberg equilibrium test at p=1*10^−9^. We further set a minor allele frequency threshold of 0.001 leaving 8,208,114 SNPs. The GWAS on each pre-residualized and normalized regional brain morphology measures was carried out using the standard additive model of linear association between genotype vector, *g_j_*, and phenotype vector, y, using PLINK2^15^. The summary statistics were subsequently formatted according to LDSC standards (https://github.com/bulik/ldsc/wiki/Summary-Statistics-File-Format).

### MiXeR analysis

We applied causal mixture models^3,9^ to the GWAS summary statistics, using the MiXer tool (https://github.com/precimed/mixer). For each SNP, *i*, univariate MiXeR models its additive genetic effect of allele substitution, *β_i_*, as a point-normal mixture, 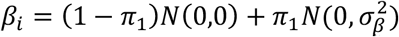, where *π*_1_ represents the proportion of non-null SNPs (‘polygenicity’) and 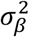 represents variance of effect sizes of non-null SNPs (‘discoverability’). Then, for each SNP, *j*, MiXeR incorporates LD information and allele frequencies for M=9,997,231 SNPs extracted from 1000 Genomes Phase3 data to estimate the expected probability distribution of the signed test statistic, 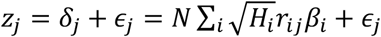, where *N* is sample size, *H_i_* indicates heterozygosity of i-th SNP, *r_ij_* indicates allelic correlation between i-th and j-th SNPs, and 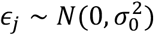 is the residual variance. Further, the three parameters, 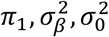, are fitted by direct maximization of the likelihood function.

The number of causal variants is estimated as *Mπ*_1_, where M=9,997,231 gives the number of SNPs in the reference panel. Phenotypic variance explained on average by a causal genetic variant is calculated as 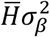, where 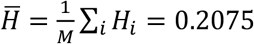 is the average heterozygosity across SNPs in the reference panel. Under the assumptions of the MiXeR model, SNP-heritability can be calculated as 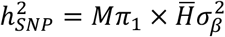.

In the cross-trait analysis, MiXeR models additive genetic effects as a mixture of four components, representing null SNPs in both traits (*π*_0_); SNPs with a specific effect on the first and on the second trait (*π*_1_ and *π*_2_, respectively); and SNPs with non-zero effect on both traits (*π*_12_). In the last component, MiXeR models variance-covariancer matrix as 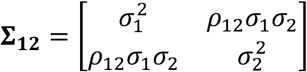 where *ρ*_12_ indicates correlation of effect sizes, and 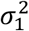 and 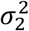 correspond to the discoverability parameter estimated in the univariate analysis of the two traits. After fitting parameters of the model, the Dice coefficient of polygenic overlap is then calculated as 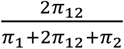, and genetic correlation is calculated as 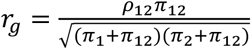.

After obtaining MiXeR parameter estimates, we excluded from our analyses regions where the ratio between the estimated heritability and its standard error (SE) was less than 3. This was done as it indicates the GWAS was insufficiently powered to reliably estimate MiXer parameters^9^

## Extended Data

### Relation of brain region size with polygenicity, discoverability and heritability

Previous studies have indicated that brain region size23 correlates with heritability estimates. We therefore investigated the relationship of region size, quantified as mean surface area, with polygenicity, discoverability and heritability. We used Spearman’s rank correlation to test for significant associations between region size and the parameters obtained from the GWAS on regional cortical surface area and thickness separate. Please see Table S1 for the full results. As reported in the main text, region size was significantly associated with all three parameters obtained from the summary statistics of regional surface area, while none were significantly associated with the regional cortical thickness-based estimates.

**Table S1.** Results from Spearman rank correlation tests between brain region size and MiXer parameter estimates, split by regional surface area and thickness-based estimates.

|  | Surface area |  | Thickness |  |
| --- | --- | --- | --- | --- |
|  | R<sub>s</sub> | p-value | R<sub>s</sub> | p-value |
| NC | -.254 | .047 | .110 | .397 |
| $\sigma^2_{\beta}$ | .412 | .001 | .022 | .866 |
| h<sup>2</sup> | .387 | .002 | .216 | .094 |

### Parameter estimates per region

Please see Figure S1 for a visual representation of the polygenicity, discoverability and heritability of each of the regional cortical measures derived from the Desikan-Killiany atlas.

### Comparisons of cortical thickness-based parameter estimates between parcellation schemes

In the main text, the comparison between the different parcellation schemes reflects regional surface area-based estimates only. In Figure S2, we present the results from the same comparisons based on cortical thickness. As mentioned in the main text, and as can be seen in the figure, there were no differences in discoverability of cortical thickness between the schemes, and only one significant difference in polygenicity, between Yeo-17 and the Glasser parcellation. Note that the Chen *et al*. parcellation is missing from this comparison. None of the 24 clusters passed the inclusion criterion of having a heritability estimate that is three times higher than its standard error. This attests to the specificity of this scheme, i.e. clusters formed on the basis of surface area do not produce meaningful cortical thickness estimates.

**Figure S1.**
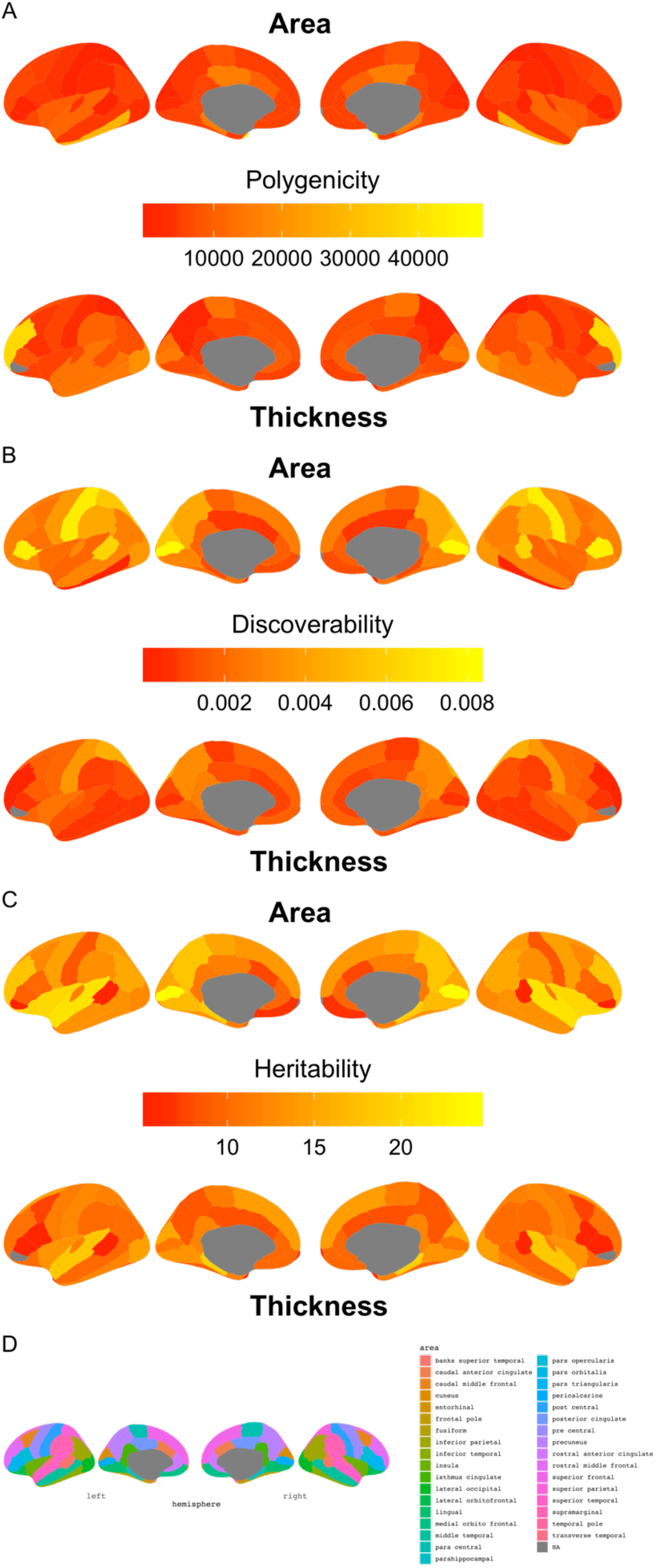
Brain maps with color coding reflecting MiXer parameter estimates for the Desikan-Killiany regional cortical measures, with A) indicating polygenicity, B) discoverability, and c) heritability. For each subplot, the top row indicates the estimates obtained from the regional surface area measures, and the bottom row those based on the regional thickness estimates. D) provides a legend o f the regions.

**Figure S2.**
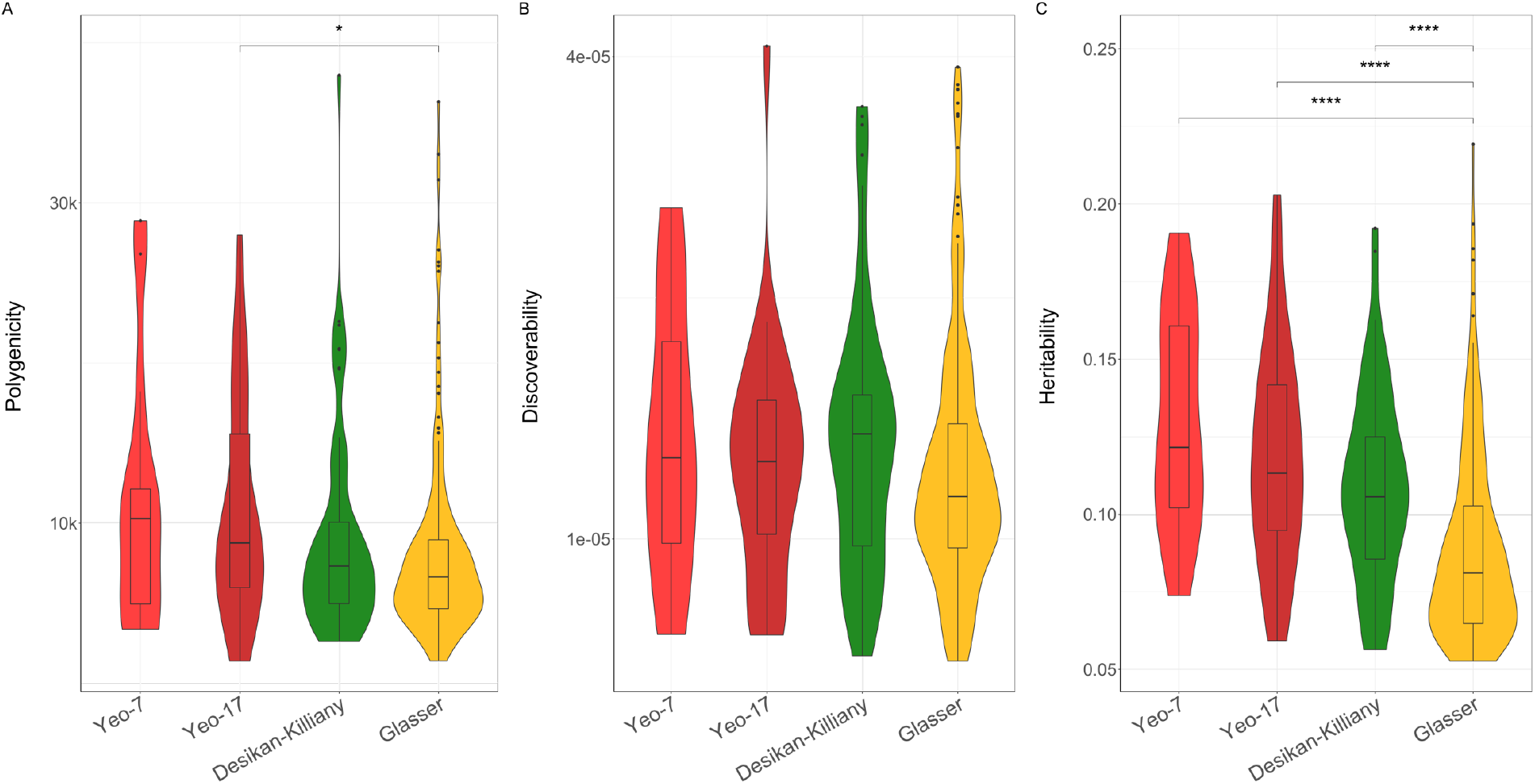
Comparisons of discoverability and polygenicity of regional cortical thickness across parcellation schemes. Violin plots comparing the polygenicity (A), discoverability (B) and heritability (C) (on the y-axis) of the different parcellation schemes (x-axis). Significance of each paired comparison, indicated at the top of each graph, is calculated through the Wilcoxon signed-rank test with *:p<.05 and ****:p<.00001.

## References

1. Grasby, K. L. et al. The genetic architecture of the human cerebral cortex. bioRxiv 399402 (2018). doi:10.1101/399402

2. Fan, C. C. et al. Beyond heritability: Improving discoverability in imaging genetics. Hum. Mol. Genet. (2018). doi:10.1093/hmg/ddy082

3. Holland, D. et al. Beyond SNP Heritability: Polygenicity and Discoverability of Phenotypes Estimated with a Univariate Gaussian Mixture Model. bioRxiv 498550 (2019). doi:10.1101/498550

4. Hogstrom, L. J., Westlye, L. T., Walhovd, K. B. & Fjell, A. M. The structure of the cerebral cortex across adult life: age-related patterns of surface area, thickness, and gyrification. Cereb. cortex 23, 2521–2530 (2012).

5. Schnack, H. G. et al. Changes in Thickness and Surface Area of the Human Cortex and Their Relationship with Intelligence. Cereb. Cortex 25, 1608–1617 (2015).

6. Panizzon, M. S. et al. Distinct genetic influences on cortical surface area and cortical thickness. Cereb. Cortex bhp026 (2009).

7. Winkler, A. M. et al. Cortical thickness or grey matter volume? The importance of selecting the phenotype for imaging genetics studies. Neuroimage 53, 1135–1146 (2010).

8. Smeland, O. B. et al. Discovery of shared genomic loci using the conditional false discovery rate approach. Hum. Genet. 1–10 (2019).

9. Frei, O. et al. Bivariate causal mixture model quantifies polygenic overlap between complex traits beyond genetic correlation. Nat. Commun. 10, 2417 (2019).

10. Andreassen, O. A., Thompson, W. K. & Dale, A. M. Boosting the power of schizophrenia genetics by leveraging new statistical tools. Schizophr. Bull. 40, 13–17 (2013).

11. Sudlow, C. et al. UK biobank: an open access resource for identifying the causes of a wide range of complex diseases of middle and old age. PLoS Med. 12, e1001779 (2015).

12. Miller, K. L. et al. Multimodal population brain imaging in the UK Biobank prospective epidemiological study. Nat. Neurosci. 19, 1523–1536 (2016).

13. Desikan, R. S. et al. An automated labeling system for subdividing the human cerebral cortex on MRI scans into gyral based regions of interest. Neuroimage 31, 968–980 (2006).

14. Rosen, A. F. G. et al. Quantitative assessment of structural image quality. Neuroimage 169, 407–418 (2018).

15. Chang, C. C. et al. Second-generation PLINK: rising to the challenge of larger and richer datasets. Gigascience 4, 7 (2015).

16. Bulik-Sullivan, B. et al. An atlas of genetic correlations across human diseases and traits. Nat. Genet. 47, 1236 (2015).

17. Chen, C.-H. et al. Hierarchical genetic organization of human cortical surface area. Science 335, 1634–1636 (2012).

18. Yeo, B. T. T. et al. The organization of the human cerebral cortex estimated by intrinsic functional connectivity. J. Neurophysiol. 106, 1125–1165 (2011).

19. Glasser, M. F. et al. A multi-modal parcellation of human cerebral cortex. Nature 536, 171 (2016).

20. Watanabe, K. et al. A global overview of pleiotropy and genetic architecture in complex traits. Nat. Genet. 1–10 (2019).

21. Bansal, V. et al. Genome-wide association study results for educational attainment aid in identifying genetic heterogeneity of schizophrenia. bioRxiv 114405 (2018). doi:10.1101/114405

22. Le Hellard, S. et al. Identification of gene loci that overlap between schizophrenia and educational attainment. Schizophr. Bull. 43, 654–664 (2016).

23. Eyler, L. T. et al. A comparison of heritability maps of cortical surface area and thickness and the influence of adjustment for whole brain measures: a magnetic resonance imaging twin study. Twin Res. Hum. Genet. 15, 304–314 (2012).

24. Patel, S. et al. Heritability estimates of cortical anatomy: the influence and reliability of different estimation strategies. Neuroimage 178, 78–91 (2018).

25. Bycroft, C. et al. Genome-wide genetic data on~ 500,000 UK Biobank participants. bioRxiv 166298 (2017).

